# YFR016c/Aip5 is part of an actin nucleation complex in budding yeast cells

**DOI:** 10.1101/610980

**Authors:** Oliver Glomb, Lara Bareis, Nils Johnsson

## Abstract

The polarisome comprises a network of proteins that organizes polar growth in yeast and filamentous fungi. The yeast *Saccharomyces cerevisiae* formin Bni1 and the actin-nucleation-promoting factor Bud6 are subunits of the polarisome that together catalyse the formation of actin filaments below the tip of budding yeast cells. We identified YFR016c (Aip5) as interaction partner of Bud6 and the polarisome scaffold Spa2. Yeast cells lacking Aip5 display a reduced number of actin cables. Aip5 binds with its N-terminal region to Spa2 and with its C-terminal region to Bud6. Both interactions collaborate to localize Aip5 at bud tip and neck, and are required to stimulate the formation of actin cables. Our experiments characterize Aip5 as a novel subunit of a complex that regulates the number of actin filaments at sites of polar growth.

**Summary statement:** YFR016c/Aip5 binds to the polarisome components Bud6 and Spa2 and supports the polarisome in the formation of actin filaments in yeast cells.

## Introduction

The polarisome is required for tip growth in budding yeast and acts as a central hub for coordinating exocytosis with actin cable formation(Tcheperegine, Gao & Bi, 2005, Sheu et al., 1998, Zahner, Harkins & Pringle, 1996, Evangelista et al., 1997). It consists of the central scaffold proteins Spa2 and Pea2, of the actin nucleation-promoting factor Bud6, the yeast formin Bni1, and of Msb3/4, GTPase activating proteins for the Rab GTPase Sec4(Sheu et al., 1998, Tcheperegine, Gao & Bi, 2005, Fujiwara et al., 1998, Evangelista et al., 1997).

Fluorescent labelling of any of the six core components results in a focussed signal at the bud tip upon bud emergence that vanishes during isotropic growth and reappears at the bud neck upon the onset of cytokinesis (Neller et al., 2015). A deletion of any of the core constituents results in a decreased intensity and less focussed fluorescent signal of all other members. These mutants display a visible defect in tip growth leading to a rounder cell shape of the buds (Tcheperegine, Gao & Bi, 2005).

The polarisome subunits Bni1 and Bud6 form a complex that nucleates actin cables (Moseley, Goode, 2005, Graziano et al., 2011). Bud6 contains a C-terminal core domain that binds Bni1, followed by a “WH2-like” domain that binds G-Actin (Graziano et al., 2011, Park et al., 2015). Binding to Bud6 favours the open, active conformation of Bni1 that accepts G-actin from Bud6 to nucleate and elongate the actin filament. The functions of the N-terminal region of Bud6 (residues 1-364) are less well defined. It contributes to the correct localization of Bud6 and was shown to bind to the microtubule (+) end binding protein Bim1 (Jin, Amberg, 2000, Ten Hoopen et al., 2012).

The polarisome interacts with a variable set of factors and changes its composition during the cell cycle (Moreno et al., 2013). Understanding the multiple functions of the polarisome thus requires a complete list of its associated factors. The protein of the yeast ORF YFR016c (Aip5) was shown to co-precipitate with Spa2 and was enriched in Bni1- and Las17-induced actin networks (Shih et al., 2005, Miao et al., 2013, Michelot et al., 2010). Aip5 is a large protein of 1233 amino acids that contains as recognizable sequence features only a glutharedoxin (GRX)-like domain at its very C-terminus. The GRX-like domain misses the consensus sequence CXXC/S that is required for catalytic activity (Fernandes, Holmgren, 2004, Holmgren, 1989). We show that Aip5 stimulates the nucleation of actin filaments by directly interacting with Spa2 and Bud6.

## Results

### Aip5 interacts with the polarisome components Spa2 and Bud6

As part of our effort to map the protein interaction network of the polarity proteins in yeast we searched for interaction partners of Aip5 by a systematic Split Ubiquitin interaction screen. Aip5 was tested as CRU fusion (Aip5CRU, **C**-terminal Ubiquitin-**R** (arginine)-**U**ra3) against an array of 533 yeast strains each expressing a different N_ub_ fusion (N-terminal half of Ubiquitin). In addition to Spa2, the assay detected Bud6 as a novel interaction partner of Aip5 (Fig. 1A, Fig. S1 for a complete list). Split-Ub analysis of Aip5CRU in cells lacking Bud6 or Spa2 demonstrated that both polarisome components interact independently from each other with Aip5 (Fig. 1B).

**Figure 1.**
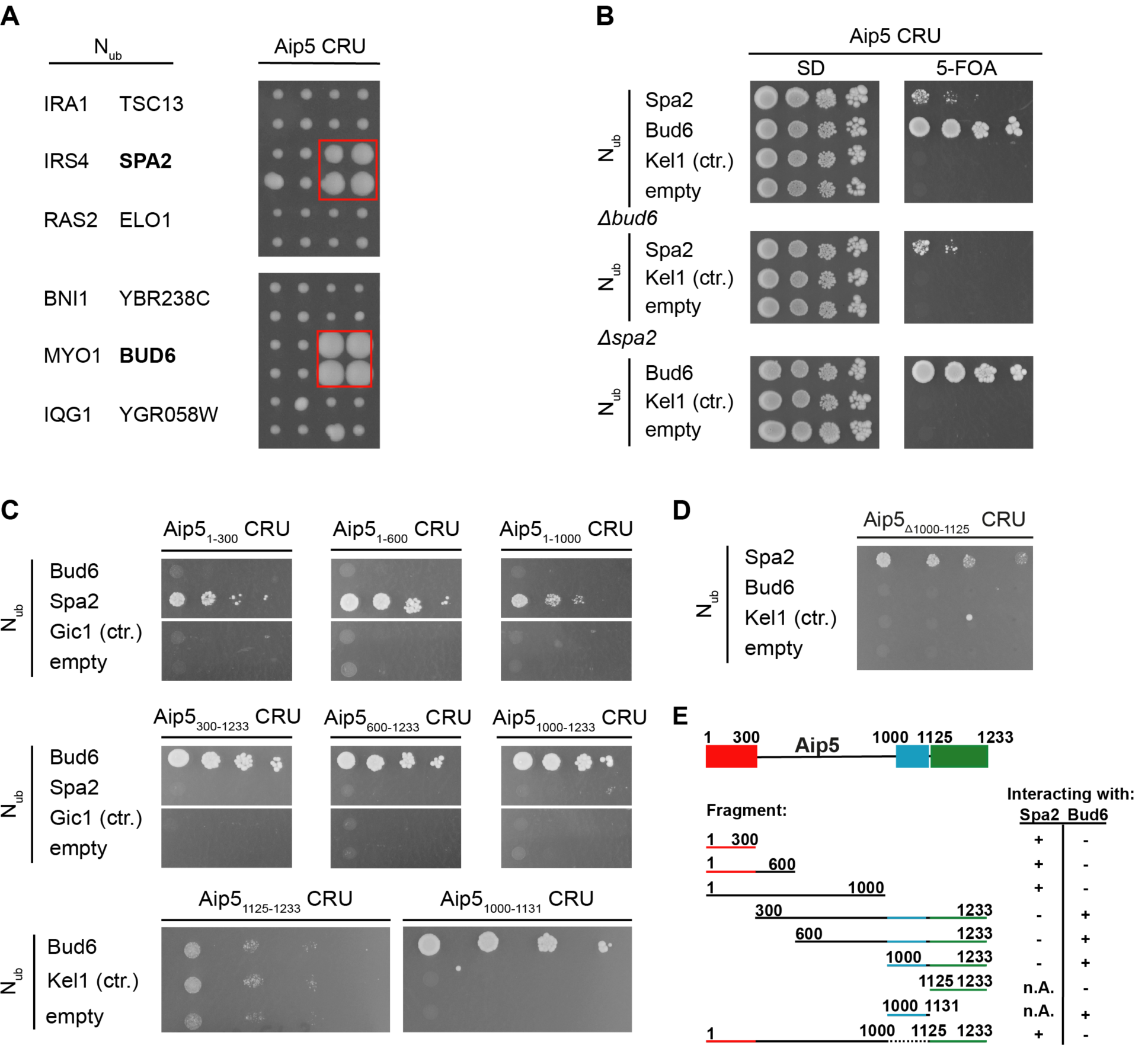
Aip5 interacts with Bud6 and Spa2 *in vivo*. (A) Right panel shows the cut outs of a split -Ubiquitin analysis of 533 diploid Aip5 CRU-containing yeast strains, each co-expressing a different N_ub_ fusion protein. The N_ub_ and CRU-expressing cells were independently mated four times, spotted in quadruplets, transferred onto medium containing 5-FOA and incubated for four days. Growth of four colonies indicates interaction. Left panel identifies the N_ub_ fusions of the corresponding cut outs. N_ub_-Bud6- and Nub-Spa2 expressing cells are boxed (see also Figure S1 for the complete array). (B) Bud6 and Spa2 interact independently of each other with Aip5. Haploid yeast strains (upper panel) lacking either *BUD6* (middle panel) or *SPA2* (lower panel) and expressing the indicated N_ub_ fusions were spotted in 10-fold serial dilutions starting at OD_600_= 1 on medium containing (right panel) or not containing (left panel) 5-FOA. Cells were incubated for seven days. (C) Split-Ub analysis as in (B) but with diploid yeast cells expressing full length Aip5 or fragments of Aip5 as CRU fusions together with the indicated N_ub_ fusions. (D) Split-Ub assay as in (B) but with diploid yeast cells expressing genomically integrated Aip5_Δ1000-1125_ CRU and co-expressing the indicated N_ub_ fusions. (E) Summary of the binding-site analysis. Spa2 binding site (red), Bud6 binding site (blue) and the predicted Grx-like domain (green) are highlighted in colours.

We next mapped the interaction sites for Spa2 and Bud6 on Aip5 by testing N- and C-terminal truncations of Aip5CRU against the N_ub_ fusions to Spa2 and to Bud6 (Fig. 1C). The assay revealed that the N-terminal 300 amino acids of Aip5 were sufficient to maintain the interaction with Spa2, whereas a deletion of this fragment resulted in a loss of interaction with Spa2. In contrast, all N-terminally truncated fragments of Aip5 that contained the C-terminal amino acids 1000-1131 interacted with Bud6. Aip5_1125-1233_ covering the conserved putative GRX-like domain did not suffice to generate a strong interaction signal with Bud6 (Fig.1C, E) whereas the fragment Aip5_1000-1131_ lacking the GRX-like domain still interacted with Bud6 in the Split-Ub assay. Accordingly, full-length Aip5 lacking residues 1000-1125 (Aip5_Δ1000-1125_) interacted with Spa2 but not with Bud6 any longer (Fig. 1D, E).

Bud6 can be divided into a C-terminal region containing the binding sites to actin and Bni1 and an N-terminal region of unknown structure and with a less defined function(Jin, Amberg, 2000). By co-expressing Aip5_1000-end_CRU together with N_ub_ fusions to different fragments of Bud6, we could localize the major binding site of Bud6 for Aip5 to its N-terminal 141 residues (N_ub_-Bud6_1-141_) (Fig. 2A). To confirm that the interaction between Aip5 and Bud6 is direct, we immobilized the *E.coli-*expressed GST-Bud6_1-141_- and GST-Bud6_1-364_ onto glutathione beads. His_6_ AIP5_1000-end_ was efficiently pulled down by GST-Bud6_1-141_ and less efficiently by GST-Bud6_1-364_ (Fig. 2B). To provide independent evidence for the interaction between Aip5 and Spa2, we artificially relocated Spa2 to peroxisomes by fusing it to the N-terminal fragment of Pex3. By co-expressing Aip5-GFP, Aip5_1-300_-GFP, or Aip5_300-end_-GFP together with Pex3_1-45_-mCherry-Spa2 we could show that only Aip5 fusions containing an intact binding site to Spa2 relocated to the Spa2-decorated peroxisomes (Fig. 2C).

**Figure 2.**
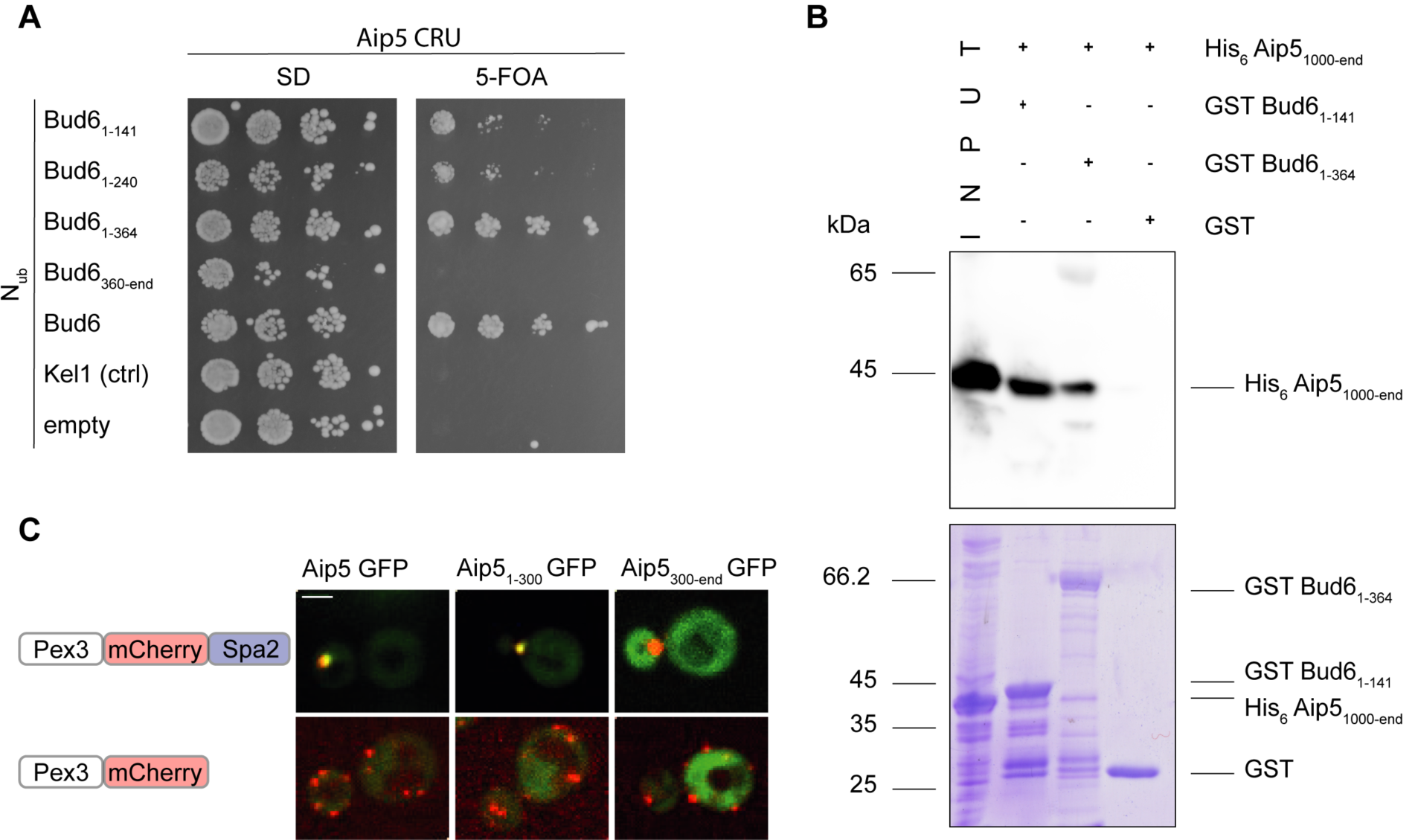
Aip5 interacts physically with Bud6 and Spa2. (A) Split-Ubiquitin analysis as in Fig. 1 (B) but with haploid yeast cells expressing Aip5_1000-end_CRU together with the indicated N_ub_-fusions. Growth on FOA indicates interaction. (B) Extracts from *E.coli* cells expressing His_6_-Aip5_1000-end_ (lane1, „input”) were incubated with glutathione-coupled beads displaying GST-Bud6_1-141_ (lane2), or GST-Bud6_1-364_ (lane 3), or GST (lane 4). Upon elution, proteins were separated by SDS-PAGE, and either stained with coomassie brilliant blue (lower picture) or with an α-His antibody after transfer onto nitrocellulose (upper picture). (C) Reconstitution of the Aip5-Spa2 interaction at the peroxisome. Upper panel: Co-localization of Aip5-GFP and selected fragments of Aip5-GFP (green) with Spa2-mCherry decorated peroxisomes (red) in cells carrying a deletion of the genomic *SPA2*. Lower panel: as above but with cells not expressing Spa2-mCherry on peroxisomes. All images are z-stack projections of the maximum intensity of 10 stacks. Scale bar indicates 2µm.

### Spa2 localizes Aip5 to sites of polar growth

A fluorescently tagged Aip5 matches the cell cycle dependent localization of a typical polarisome component. AIP5-GFP assembled at the incipient bud site in late G1, stayed focused at the cell tip during bud growth, and re-located to the bud neck during cytokinesis (Fig. 3A). To determine whether the localization of Aip5 depends on either Bud6 or Spa2, we compared the fluorescence intensities of Aip5-GFP between wild type-, *Δspa2*-, or *Δbud6*-cells (Fig.3 B, C). The localization of Aip5 significantly decreased at bud tip and bud neck in both *Δbud6*- and *Δspa2* cells. We conclude that both polarisome subunits collaborate to localize Aip5 at sites of polar growth. The major determinant of its cellular localization is however provided by the interaction with Spa2. The cytosolic distributions of GFP fusions to different fragments of Aip5 showed that only AIP5_1-300_-GFP, containing the binding site for Spa2, was still measurably enriched at the bud (Fig. 3D). Aip5 does not contribute to the localization of Spa2 or Bud6 (Fig. S2).

**Figure 3.**
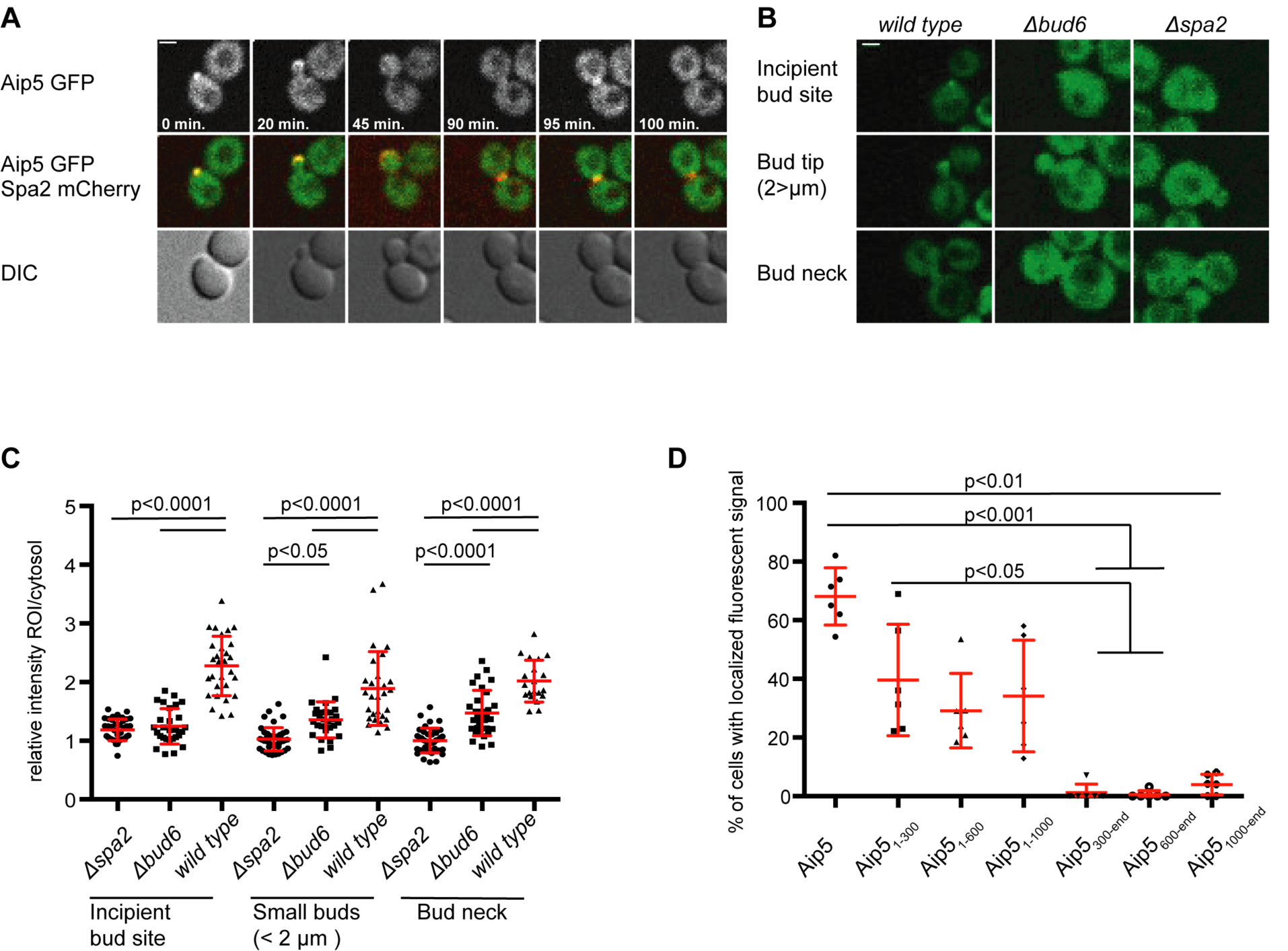
Spa2 and Bud6 attract Aip5 to sites of polar growth. (A) Time-resolved localization of Aip5-GFP (upper and middle panel) and Spa2-mCherry (red, middle panel) over one cell cycle. All images are derived from single planes from a confocal microscope. DIC images are shown in the lower panel. (B) Wild type-, *Δspa2-* and *Δbud6-*cells expressing Aip5-GFP were analysed by fluorescence microscopy. A single plane of the confocal image was used to calculate the mean fluorescence at the indicated positions and normalized to the cytosol of the mother cell. Error bars indicate the standard deviations. (C) Quantitative analysis of the cells shown in (B). Statistical analysis was performed with a one-way Anova followed by Bonferroni’s post-test with n*(Δspa2*_Incipient Bud Site (IBS)_)=38, n*(Δbud6*_IBS_)=28, n(wild type_IBS_)=29, n(*Δspa2*_Small_ Buds(SB))=44, n(*Δbud6*_SB_)=28, n(wild type_SB_)=29, n(*Δspa2*_Bud Neck (BN)_)=29 n(*Δbud6*_BN_)=40, n(wild type_BN_)=30 cells of two independent experiments. (D) Yeast cells expressing Aip5-GFP (n=207) or the GFP-labelled Aip5-fragments Aip5_1-_ 300 (n=239), Aip5_1-600_ (n=213), Aip5_1-1000_ (n=191), Aip5_300-end_ (n=165), Aip5_600-_ end (n=143) and Aip5_1000-end_ (n=188) were analysed by fluorescence microscopy. Images were taken and z-stack projections of the maximum intensity were used to calculate the percentage of cells (containing buds < 2µm) with a polarized fluorescence signal. Mean values of two single transformants for each fragment were pooled and corresponding standard deviations from three independent measurements are shown. Statistical analysis was performed with a Kruskal-Wallis test followed by Dunn’s multiple comparison. Mean values with standard deviations are shown in red.

### Aip5-Bud6 increase the cellular resistance against latrunculin A

Bud6 and Aip5 were found in a genome-wide screen for genes that influence the sensitivity for the actin depolymerizing agent latrunculin A (LatA) (Hoepfner et al., 2014).

To test whether Aip5 and Bud6 act in the same pathway, we compared the extent of LatA growth inhibition between wildtype-, *Δaip5-, Δbud6-* and *Δaip5Δbud6-*cells. The single deletion of *AIP5* conferred a higher LatA sensitivity than the deletion of *BUD6* whereas the additional deletion of *BUD6* in a *Δaip5* strain did not further increase the LatA sensitivity of the strain (Fig. 4A). Accordingly, a fragment of Aip5 harbouring the Bud6-binding region and the conserved GRX-like domain sufficed to reduce the LatA sensitivity of *Δaip5* cells (Fig. 4B). Neither the isolated Bud6-binding site nor the C-terminally located GRX-like domain complemented the LatA sensitivity of *Δaip5* cells (Fig.4C).

**Figure 4.**
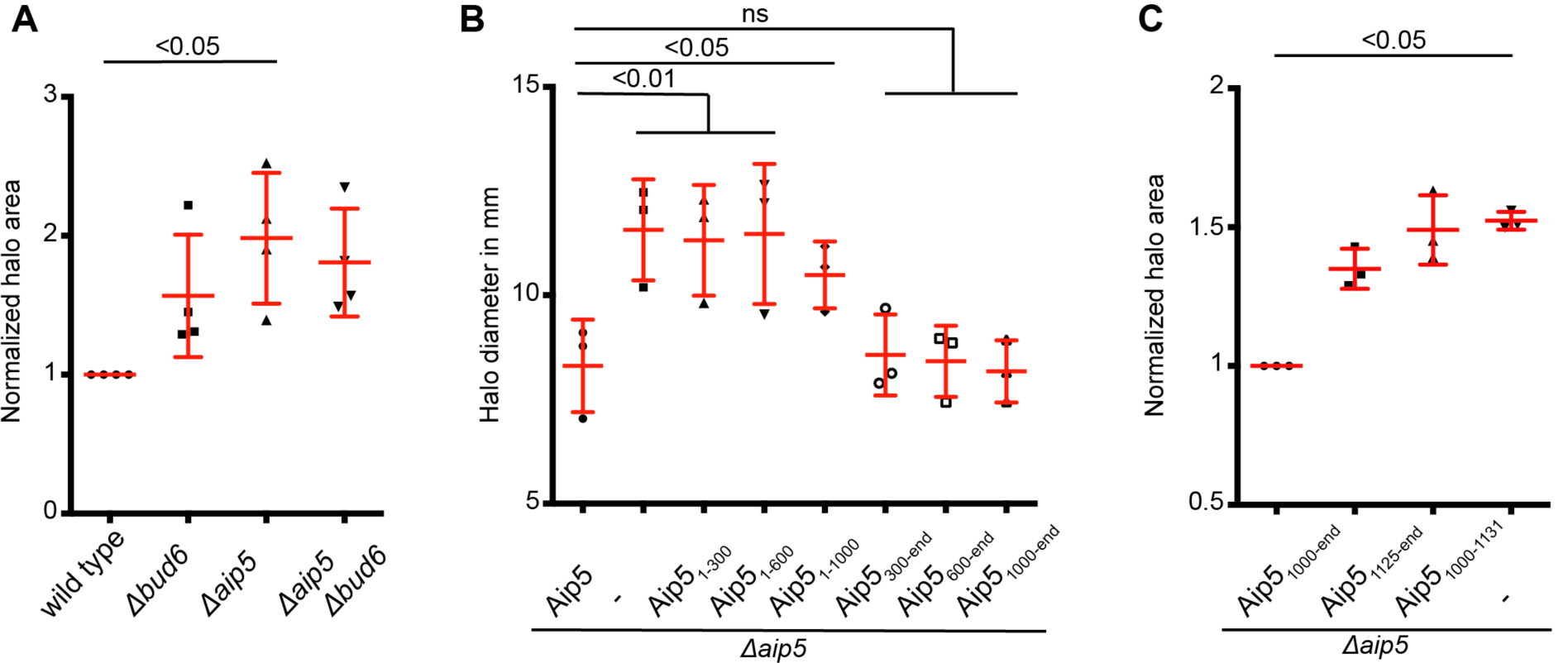
A fragment of Aip5 including the Bud6 binding site and the GRX-like domain rescues the LatA sensitivity of *Δaip5*-cells. (A) Wild type-, *Δaip5-, Δbud6-* and *Δaip5Δbud6-*cells were incubated on SD plates in the presence of a centrally placed filter disk soaked with 10 µl LatA (500 µM). Values of the normalized area of growth inhibition were collected from four independent measurements (n=4) and compared by Kruskal-Wallice test and a Dunns post test. (B) *Δaip5* cells expressing different fragments of Aip5 CRU were incubated as in (A) in the presence of 250 µM LatA. The diameter of the zone of growth inhibition was measured and compared by one-way ANOVA and a Tukeys post test. Values were collected from three independent measurements (n=3). (C) *Δaip5*-cells expressing the indicated fragments of Aip5 as CRU fusions or an empty control plasmid were incubated as in (A) but on SD plates lacking methionine in the presence of 500 µM LatA. Values of the normalized area of growth inhibition were collected from three independent measurements (n=3) and compared by Kruskal-Wallis test and a Dunns post test. Mean values with standard deviations are shown in red.

### Aip5 supports Bud6-induced actin filament formation

The LatA sensitivities of *AIP5*-alleles lacking the Bud6-binding site suggest that Aip5 might support actin filament nucleation by Bud6. We performed actin-phalloidin staining to compare the number of actin filaments between wild type-, *Δaip5, Δbud6* and *Δaip5Δbud6*-cells. The average of four actin filaments was reduced to three in *Δaip5* and to 1.5 filaments in *Δbud6* cells. Deleting *AIP5* in *Δbud6*-cells did not further reduce this number (Fig. 5 A).

**Figure 5.**
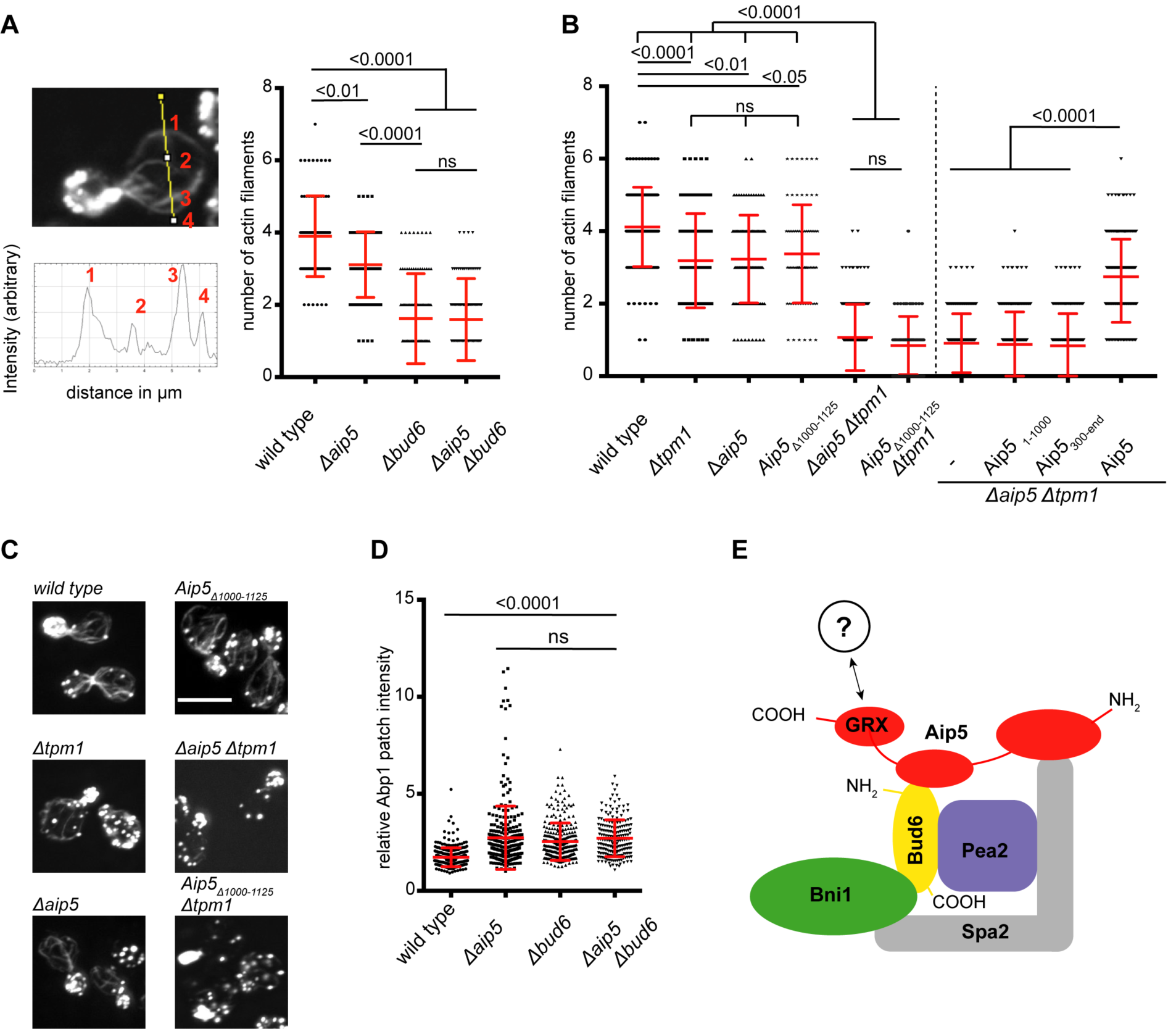
Aip5 supports actin filament formation by Bud6. (A) Left panel: Wild type-(n=98), *Δaip5-* (n=98), *Δbud6-* (n=94) and *Δaip5Δbud6-*cells (n=93) were fixed in PFA and their actin was stained with Alexa488-Phalloidin. Intensity plot profiles were calculated in budding mother cells perpendicular to the mother-bud axis. The local maxima were counted to determine the number of actin filaments in this section. Right panel: Filament numbers were compared among cells of the indicated genotypes using a Kruskal-Wallis test and a Dunns post test. Error bars show the standard deviations of three independent measurements. (B) The average actin filament number was determined as in (A) for the indicated genotypes in two different experimental setups divided by the vertical dashed line. Left side: Wild type- (n=138), *Δtpm1*- (n=140), *Δaip5-* (n=100), Aip5_Δ1000-1125_- (n=75), *Δtpm1Δaip5-* (n=192) and *Δtpm1*, Aip5_Δ1000-1125_-cells (n=71) were compared for their average actin filament number by a Kruskal-Wallis test followed by a Dunns post test. The values were taken from two independent measurements. Right side: *Δtpm1Δaip5* cells were transformed with an empty plasmid (control, n=222), or with plasmids expressing Aip5_1-1000_ (n=258), Aip5_300-end_ (n=195) and Aip5 (n=255). The average filament numbers pooled from two independent measurements were compared by Kruskal-Wallis- and a Dunns post-test. (C) Representative images of actin stainings from cells shown in (B, left side). Images are maximum intensity z-projections of 14 single z-stacks taken with a confocal microscope. (D) The mean fluorescence intensities of Abp1-GFP-decorated actin patches were normalized to the mean fluorescence intensity of cytosolic Abp1-GFP and compared between yeast cells of the wild type (n=298), *Δaip5-* (n=289), *Δbud6-* (n=286) and *Δaip5Δbud6-*cells (n=201). Each value represents a single Abp1 patch. Measurements were performed in triplicates. Error bars indicate standard deviations. Abp1 intensities were compared among genotypes by a Kruskal-Wallis-test and Dunns post test. (E) Proposed model of the actin filament nucleation complex. See discussion for details. Mean values with standard deviations are shown in red.

The yeast tropomyosin Tpm1 decorates and thereby stabilizes the once formed linear actin filaments (Drees et al., 1995). By removing Tpm1, we wished to sensitize the cells for modest changes in their ability to nucleate new actin filaments. The average number of actin cables was reduced to three in *Δtpm1* cells (Fig. 5B, C). The simultaneous deletion of *AIP5* further reduced this number to one. Only the full length Aip5 reverted the decrease of actin filaments in *Δtpm1Δaip5*-cells. N- or C-terminally truncated fragments of Aip5 lacking the binding site for Spa2 or the Bud6 binding site and the GRX-like domain did not stimulate actin filament formation (Fig. 5B). Aip5_Δ1000-1125_ lacking only the binding site to Bud6 also failed to substitute for Aip5 in this assay (Fig. 5B).

Actin patches consist of cross-linked actin networks that support endocytosis in the bud of yeast cells(Kaksonen, M., Sun & Drubin, 2003, Kaksonen, Marko, Toret & Drubin, 2005, Munn, 2001). Abp1 is a late endocytotic protein that interacts with actin in these structures(Kaksonen, M., Sun & Drubin, 2003). To test the contribution of Aip5 to actin patch formation, we expressed Abp1-GFP in wild type-, *Δaip5-, Δbud6-* and *Δaip5Δbud6-*cells and compared the relative GFP-intensities of the actin patches between these cells. A deletion of *BUD6* or *AIP5* enhanced the signal intensity of fluorescently labelled Abp1 in actin patches (Fig. 5D). The double deletion of *AIP5* and *BUD6* did not amplify the effect of the single deletions (Fig. 5D). We conclude that Aip5 is not involved in actin patch formation and assume that its effect on actin patch intensity is an indirect consequence of a larger pool of free G-actin in these cells (Shin et al., 2018).

## Discussion

Originally described as a protein that co-precipitates with the polarisome core component Spa2, and found associated with actin structures we could now demonstrate by actin staining of intact yeast cells that Aip5 is involved in the formation of actin filaments (Shih et al., 2005; Miao et al., 2013). This activity is the collaborative action of a least two binding sites of Aip5. The N-terminal 300 residues of Aip5 harbour the binding site for Spa2. The second site lies between residues 1000 and 1131 and binds directly to Bud6 (Fig. 5E). Mutants of Aip5 that lack either of the two binding sites display a smaller number of actin filaments in otherwise wildtype- or *Δtpm1*-cells. The Spa2-binding site is the major determinant for the localization of Aip5 to bud tip and bud neck. As both locations are the sites of the highest filament nucleation activity, we propose that Spa2 is required to anchor Aip5 to these sites and brings it into close proximity of the actin nucleation/elongation factors Bud6 and Bni1 (Fig. 5E). The Bud6-binding site contributes to the correct localization of Aip5 but has a second function that is more directly involved in actin turnover. Cells lacking Aip5 are more sensitive to latrunculin A, and the Bud6-binding site of Aip5 is required to complement this sensitivity. LatA reduces the amount of polymerizable G-actin(Ayscough et al., 1997). Consequently, the lack of factors that stimulate G-actin to form linear actin filaments should make the cell more sensitive for LatA. The direct binding to Bud6 suggests that Aip5 exerts its stimulating activity through activating Bud6. This assumption is supported by the observation that similar to Aip5 the deletion of Bud6 leads to less actin filaments and a higher LatA sensitivity. The simultaneous deletion of both genes is not additive with respect to the strength of both phenotypes. This, together with the found physical interaction, suggests that Aip5 and Bud6 work in a complex to stimulate the formation of actin filaments. How binding to the N-terminal domain might change the C-terminally located activity of Bud6 is not known and will require further structural work.

Aip5 harbours at its very C-terminus a GRX-like domain (Fig. 5E). Together with the neighbouring Bud6-binding site, the GRX-like domain is required for the rescue of the LatA sensitivity of *Δaip5* cells. We can only speculate about the activity of this domain, but binding to Bni1, G-actin or profilin would fit well with the proposed activities of Aip5 (Fig. 5E) (Moseley et al., 2004, Evangelista et al., 2002, Imamura et al., 1997). This idea is supported by the synthetic sickness of cells carrying a deletion of *AIP5* and certain alleles of profilin that impair either actin- or formin binding (Pfy1-4 and Pfy1-13) (Wolven et al., 2000, Costanzo et al., 2016).

## Materials and Methods

All yeast strains used in this study are derivatives of the *Saccharomyces cerevisiae* strain JD47 (Dohmen et al., 1995) (Table S1). Cells were grown at 30°C in YPD or synthetic complete (SD) medium lacking certain amino acids or containing the indicated antibiotics to select for the presence of gene fusion constructs. For co-localization analysis of cells containing Spa2-decorated peroxisomes, the o.n. culture was diluted into medium lacking methionine to enhance the expression of the Pex3-Spa2 chimera. *E.coli* strain XL1 blue was used for plasmid preparation. *E.coli* strain BL21 DE3 was used for protein production. Both *E.coli* strains were grown at 37°C in LB medium with appropriate antibiotics added.

### Construction of plasmids and strains

Fusions of GFP and CRU to full-length *AIP5* or fragments of it, were constructed by PCR amplification from yeast genomic DNA with appropriate primers containing *EagI* or *SalI* restriction sites(Wittke et al., 1999). The amplified fragments were digested with *EagI/SalI* and ligated in front of a GFP/CRU module in the vector pRS313 (Sikorski, Hieter, 1989). The expression of Aip5 fragments was controlled from a P_*MET17*_ promoter. Plasmids containing N_ub_ Bud6 fragments were cloned in frame into a pCUP1*-*Nub-HA kanMX4, Cen plasmid behind the N_ub_ module with primers containing *SalI/Acc65I* restriction sites.

For the construction of the genomically integrated CRU and GFP fusions a PCR fragment spanning the C-terminal ORF of *AIP5* using the primer „YFR016c Cub Eag” and „YFR016c Cub SAL” without stop codon and containing *Eag*I and a *Sal*I restriction sites was cloned in front of the CRU module in a pRS303 vector or in front of GFP in a pRS304 vector (Sikorski, Hieter, 1989). For the integration into the *AIP5* locus the plasmid p*AIP5CRU303* was linearized using a single *AflII* site in the genomic *AIP5* sequence. Successful integration was verified by PCR of single yeast colonies with the diagnostic primer combination (5’->3’) „YFR016c Cub Ctr 450” and „Cub Ctr”. The plasmid *pAIP5GFP304* was linearized using a single *BglII* site in the genomic *AIP5* sequence. Successful integration was verified by PCR of single yeast colonies with the diagnostic primer combination (5’->3’) „YFR016c Cub Ctr 450” and „GFP (S65T) ATG+84RV”. Gene deletions were obtained by replacing the ORF with an antibiotic resistance cassette through single step homologous recombination as described (Janke et al., 2004). Briefly, resistance cassettes were amplified by PCR using primers carrying sequences at their 5’end identical to stretches directly up-/downstream to the gene of interest (S1, S2). Successful gene deletion was verified by single colony-PCR using primers up/downstream of the ORF (G1, G2) as well as primers annealing within the resistance marker.

Fragments of *BUD6* or *AIP5* were expressed as GST- or His_6_-fusions in *E.coli*. GST-fusions were obtained by amplification of the respective fragments from genomic yeast DNA using primers containing *NcoI/EcoRI* restriction sites. The PCR fragment was cloned in-frame behind *GST* in the plasmid pGex6P1 or pGex2T (GE-Healthcare). The 6x His-tag was added through PCR of the respective fragment containing *Sfi*I restriction sites as well as a TEV-recognition site in the forward primer, and insertion of the resulting fragment in frame downstream of a 6x His-tag into the pAC plasmid (Schneider et al., 2013).

The chimeric Pex3_1-45_mCherry pRS306 plasmid was adapted from(Luo, Zhang & Guo, 2014). For the generation of the *PEX3-SPA2* chimeras a *PEX3*-fragment containing the N-terminal 45 residue of *PEX3* (Pex3_1-45_) was amplified from genomic DNA using *XbaI/EagI* restriction sites and inserted between a P_*MET17*_ promoter and mCherry in a modified vector pRS306, containing a P_*MET17*_ promoter. *SPA2* was inserted into the resulting plasmid *P*_*MET17*_*-Pex3-mCherry-mcs* in frame behind the mCherry using *BamHI/SalI* restriction sites and containing an HA tag at the 3’ end.

Guide-RNA construction into a CRISPR/CAS9 plasmid was adapted from the protocol of the Wyrick Lab (Laughery et al., 2015). gR1 AIP5-3309 and gR2 AIP5-3309 primers were designed based on this tool and hybridized at a concentration of 3 µM in 1×T4 DNA ligase buffer (NEB) by initial heating and subsequent cooling steps. The hybridized oligonucleotides were ligated into the *SwaI, BclI* restricted pML104 plasmid. For deleting residues 1000-1131 in *YFR016*, pML104-YFR016c-3309 was co-transformed with the oligonucleotide doYFRd1000-1125. Following kick-out of the pML104-YFR016c-3309 plasmid by counter selection on medium containing FOA, successful deletion of amino acids 1000-1125 was confirmed by single-colony PCR using the primers YFR016cCRISPctrl2860 and G2 YFR016c. All used plasmids and oligonucleotides are listed in Table S2 and Table S3.

### *In vitro* binding assay

PGex6P1, pGex2T and pAC expression plasmids containing GST- or 6xHis fusion proteins were transformed into *E.coli* BL21. O.n. cultures were diluted to OD_600_ 0.3 in LB or SB medium and grown to OD_600_ 0.8 at 37°C before protein expression was induced with IPTG. Following expression conditions were chosen:

His_6_ AIP5_1000-end_: 1 mM IPTG, 18°C, o.n., SB (IPTG concentration, expression temperature, -duration, -medium); GST Bud6_1-141_: 1 mM IPTG, 37°C, 4 h, LB; GST Bud6_1-364_: 1 mM IPTG, 37°C, 4 h, LB; GST: 1 mM IPTG, 37°C, 4 h, LB Cells were pelleted after incubation and stored at −80°C. Cell pellets were resuspended in 1xHBSEP (10 mM Hepes, 150 mM NaCl, 3 mM EDTA, 0.005% Tween-20) buffer containing 1× protease inhibitor cocktail (Roche Diagnostics, Penzberg, Germany) and lysed with 1 mg/ml Lysozyme for 20 min. on ice followed by a sonification for 2×4 min with a Bandelin Sonapuls HD 2070 (Reichmann Industrieservice, Hagen, Germany). Extracts were separated from cell debris by centrifugation for 10 min, 4°C at 40,000 g. Extracts containing GST, GST-Bu6_1-141_, or GST-Bud6_1-364_ were incubated with HBSEP-equilibrated glutathione-coupled sepharose beads (GE Healthcare, Freiburg, Germany), for 30 min under rotation in the cold. Labelled glutathione beads were washed twice with HBSEP and incubated with 0.1 mg/ml BSA (Sigma Chemicals, St.Louis, USA) for 30 min. Beads were finally treated with His_6_Aip5_1000-end_ extract in the presence of 0.1 mg/ml BSA for 1 h. Three washing steps with 1xHBSEP were used to remove unbound proteins and bound proteins were eluted by incubating the glutathione beads with 1× GST elution buffer (50 mM Tris, 20 mM reduced glutathione). The proteins of the eluate were separated by SDS PAGE and stained with Coomassie Brilliant Blue or with anti-His antibody after transfer onto nitrocellulose membrane (Sigma-Aldrich, Steinheim, Germany; dilution 1: 5000).

### Split-Ubiquitin analysis

Split-Ub interaction analysis was either performed against an array of 533 different N_ub_ fusion proteins by replica plate analysis or by spotting cells co-expressing a chosen pair of N_ub_ and CRU fusion proteins (for a detailed description of the procedures see(Johnsson, Varshavsky, 1994, Dunkler, Muller & Johnsson, 2012, Muller, Johnsson, 2008).

For the automated array analysis an *AIP5CRU*-expressing yeast a-strain was mated with a library of 533 different N_ub_ fusion protein-expressing yeast alpha-strains. Mating and transfer on media selecting for diploid were performed with a RoToR HDA robot as described (Singer Instruments, Somerset, UK) (Hruby et al., 2011, Dunkler, Muller & Johnsson, 2012). For manual split-ubiquitin analysis, the cells were transferred on media containing or lacking 1 mg/ml 5-fluoro-orotic acid (5-FOA, Formedium, Hunstanton, UK) and incubated at 30°C(Dunkler, Muller & Johnsson, 2012). Pictures were taken after 2 to 7 days. The matrix of all N_ub_ fusion proteins is published in (Kustermann et al., 2017, Hruby et al., 2011). The medium composition is published in(Dunkler, Muller & Johnsson, 2012).

For the manual analysis O.n.-grown cultures of haploid yeast cells each expressing a single pair of CRU and N_ub_ fusion protein were spotted in 10-fold serial dilutions starting at OD_600_=1 to 0.0001 on medium containing or lacking 1 mg/ml 5-fluoro-orotic acid (5-FOA, Formedium, Hunstanton, UK). The cells were incubated at 30°C and pictures were taken after 2 to 7 days.

### Fluorescence microscopy

Fluorescence microscopy was performed on an Axio Observer Z.1 spinning-disc confocal microscope (Zeiss, Göttingen, Germany) containing a switchable Evolve512 EMCCD (Photometrics, Tucson, USA) or an Axiocam Mrm camera (Zeiss, Göttingen, Germany). The microscope was also equipped with a Plan-Apochromat 100×/1.4 oil DIC objective and 488 nm and 561 nm diode lasers (Zeiss, Göttingen, Germany). Images were recorded with the Zen2 software (Zeiss, Göttingen, Germany) and analyzed with FIJI. Alternatively, time-lapse microscopy was performed with a DeltaVision system (GE Healthcare, Freiburg, Germany) provided with an Olympus IX71 microscope (Olympus, Hamburg, Germany). This microscope contained a CoolSNAP HQ2-ICX285 or a Cascade II 512 EMCCD camera (Photometrics, Tucson, USA), a 100× UPlanSApo 100×1.4 Oil ∞/0.17/FN26.5 objective (Olympus, Münster, Germany), a steady-state heating chamber and a Photofluor LM-75 halogen lamp (89 NORTH, ChromaTechnology, Williston, USA).

O.n. grown yeast cells were diluted to OD_600_= 0.3 and grew for 3-5 hours to exponential growth phase. 1,5 ml cells were spun down and resuspended in 30-50 µl fresh medium. 3.1 µl of this suspension were placed on a glass slide covered with a glass cover slip. For time-lapse microscopy the cells were immobilized with a glass coverslip on solid SD medium containing 1.9 % agarose on a custom-designed glass slide (Glasbläserei, Ulm University).

### Quantitative analysis of fluorescence microscopy

All microscopy files were analysed and processed using FIJI (US National Institute of Health; Version 2.0.0-rc-69) (Schindelin et al., 2012). All Images were acquired as 7 to 14 z-stack files and either single stacks or projections of single layers were analysed. To calculate the local enrichment of mean fluorescence at a certain region of interest (ROI), mean intensity of the ROI was normalized to the cytosolic mean intensity of the cell and randomly chosen areas around the cell were measured as background and subtracted from the intracellular intensities. The number of actin filaments running in parallel to the mother-bud axis were measured with the assistance of the FIJI tool „Plot Profile”. A ruler was set perpendicular to the mother-bud axis and local fluorescence intensity maxima of phalloidin-stained actin filaments were counted.

### Actin Staining

Exponentially grown cells were incubated in 3.7 % Formaldehyde for 1 h, followed by an ethanolamine (1 µM) incubation for 10 min. Following fixation, the cells were washed with PBS and incubated with 66 nM Alexa-fluorophore-coupled phalloidin (Thermo Fisher Scientific, Waltham, MA, USA) for 30 min at 4°C.

### LatA Sensitivity Assay

O.n.-grown cells were collected, resuspended in H_2_O and adjusted to OD_600_= 1. 400 µl of the cell suspension were spread on solid medium, and filter disks (5 mm diameter, Roth, Karlsruhe, Germany) soaked with 10 µl LatA of various concentrations (Sigma-Aldrich GmbH, Steinheim, Germany; dissolved in DMSO) or pure DMSO was placed on top. The cells were incubated for 1-2 days at 30°C and pictures were taken to document the extent of growth inhibition around each filter disk.

## Acknowledgements

We thank Steffi Timmermann for technical assistance and Amelie Kuhn for performing the array analysis.

